# Pairs of mutually compensatory frameshifting mutations contribute to protein evolution in vertebrates and insects

**DOI:** 10.1101/2020.12.25.424394

**Authors:** Dmitry Biba, Galya Klink, Georgii Bazykin

## Abstract

Insertions and deletions of lengths not divisible by 3 in protein-coding sequences cause frameshifts that usually induce premature stop codons and may carry a high fitness cost. However, this cost can be partially offset by a second compensatory indel restoring the reading frame. The role of such pairs of compensatory frameshifting mutations (pCFMs) in evolution has not been studied systematically. Here, we use whole-genome alignments of protein coding genes of 100 vertebrate species, and of 122 insect species, studying the prevalence of pCFMs in their divergence. We detect a total of 619 candidate pCFM-genes; 11 of them pass stringent quality filtering, including three human genes: *RAB36, ARHGAP6* and *NCR3LG1*. In some instances, amino acid substitutions closely predating or following pCFMs restored the biochemical similarity of the frameshifted segment to the ancestral amino acid sequence, possibly reducing or negating the fitness cost of the pCFM. Typically, however, the resulting sequence bore no biochemical similarity to the ancestral one, indicating that pCFMs can uncover radically novel regions of protein space. In total, pCFMs represent an appreciable and previously overlooked source of novel variation in amino acid sequences.

## Introduction

The origin of radically novel amino acid sequences remains somewhat of an enigma. Numerically, by far the most common event in evolution of protein-coding genomic segments is a single-nucleotide substitution (Murphy et al. 2004). While gradual accumulation of such substitutions can modify the characteristics, structure and, in some instances, function of the encoded protein, they only allow reaching a limited neighborhood of the existing sequence in the sequence space (Povolotskaya & Kondrashov 2010). This leaves the question of the possibility of origin of new segments of protein-coding sequences from scratch. Several mechanisms for such events are known. Some involve acquisition of new DNA segments within protein-coding regions, by repeat expansion (Hancock & Simon 2005) or non-repeat-associated insertions with lengths divisible by 3. Others involve redefinition of an existing DNA segment as protein-coding; these include emergence of exonic segments from intronic ones (Artamonova & Gelfand 2007), sequences encoding N- or C-termini of proteins from 5’-or 3’-UTRs (Vakhrusheva et al. 2011), or even entire genes from non-coding DNA (Neme & Tautz 2013), (Bornberg-Bauer et al. 2021); or acquisition of an alternative reading frame for an existing protein-coding sequence (Keese & Gibbs 1992).

Because a previously non-coding sequence is expected to carry an in-frame stop codon roughly every 20 codons, radically new protein-coding sequences are seldom long. Nevertheless, the complete novelty of the encoded sequence may result in reaching remote domains of the sequence space, which may be then fine-tuned in the course of subsequent evolution (Carvunis et al. 2012).

Another possible mechanism of origin for a new amino acid-coding sequence is frameshifting insertions and deletions (hereafter, indels). Generally, they are considered to be an unlikely source of novelty (Ohno 1970) because they usually cause major disruptions in the amino acid sequence and its truncation through gain of premature stop-codons, and therefore should be extremely rare to fix due to fitness cost associated with them. Nevertheless, there are some examples of fixation of frameshifting indels and, though the protein function is often lost in these cases, sometimes it is retained (Hahn & Lee 2005). Moreover, in some cases such indels give rise to a totally new protein function (Ohno 1984; Vandenbussche et al. 2003). Still, these mutations tend to fix only if they happen near the 5’-or 3’-end of a gene, which likely lowers their impact on protein structure (Hu & Ng 2012; MacArthur et al. 2012).

Another factor which may prevent a frameshifting indel from disrupting overly long segments of protein-coding sequence is a gain of a second compensatory indel in close proximity to the first one. If the total length of the two indels is divisible by 3, the reading frame will be restored and the only amino acids changed would be those between these indels. In experimental evolution of phages, indels are frequently compensated by frame-restoring indels -a finding which has been instrumental in the discovery of the structure of genetic code (Crick et al. 1961). However, the role of compensatory pairs of indels in natural evolution has not been studied systematically. Meanwhile, there are some reasons to believe that the changes to protein structure during frameshifts are not as substantial as replacement with a totally random sequence: some key physico-chemical properties of the encoded amino acid sequence tend to be preserved by frameshifts (Wang et al. 2016; Bartonek et al. 2020). Finally, the intermediate, yet-to-be-compensated state of a protein may not be as deleterious as one could have expected, since ribosomes often bypass premature stop-codons, which are the most detrimental feature of such state (Rockah-Shmuel et al. 2013). With all these considerations taken into account, we suggest that pairs of mutually compensatory frameshifting mutations (hereafter, pCFMs, pCFM for a single such pair) may be of importance in the evolution of novel amino acid sequences.

In this paper, we systematically study pCFMs that have occurred rapidly one after the other in the evolutionary history of vertebrate and insect genes. Using a phylogenetic reconstruction of ancestral states, we describe multiple instances of such events, study the characteristics of the proteins and protein segments in which they occur, and predict their impact on the properties of the encoded protein sequence.

## Results

### Detected pairs of compensatory frameshifting mutations

We designed a parsimony-based algorithm that infers pCFMs from a multiple sequence alignment of a gene and the corresponding phylogenetic tree (see “Inference of pCFMs” section in the Materials & Methods). Inference of frame-disrupting indels from interspecies comparisons of genomic sequences is complicated by a low signal-to-noise ratio due to sequencing, assembly or alignment errors. We focused on those pCFMs that are most likely to be reliable as follows. Firstly, we reasoned that selection against a *bona fide* frame-disrupting indel is generally expected to be strong, and therefore those indels that have survived for long periods of evolutionary time are more likely artefactual. Therefore, we considered only those pairs of indels that both occurred on the same segment of the phylogenetic tree, which means that the possible intermediate state (i.e., that carrying just one of the indels but not the other) was not observed in any extant species or internal phylogenetic nodes. Secondly, we reasoned that shorter indels are less likely to be artefactual; therefore, we only considered indels of 1 or 2 nucleotides in length. While some artefactual indels can still slip through these filters, we expect this set to be strongly enriched in true pCFMs.

Under these restrictions, we found a total of 619 genes (490 in vertebrates and 129 in insects) that carried pCFMs. We refer to this set of genes as the “low-confidence dataset”. The vast majority of these pCFMs were only observed in a single species each; multiple species inheriting the same pCFM were observed only for 15 genes in vertebrates and for 12 genes in insects (Table 1). Among them, 6 vertebrate genes and 5 insect genes (Figure 1, Table 2) were described in multiple databases and/or were supported by mRNA or protein sequence data (Supplementary Table 2; see the “Post-filtering of pCFMs” section in the Materials & Methods). We refer to these 11 genes as the “high-confidence dataset”.

**Table 1.** Number of genes surviving each of the filtering stages.

|  | Vertebrates | Insects |
| --- | --- | --- |
| Number of analysed protein-coding genes | 21208 | 15283 |
| Number of genes with pCFM comprised of indels of length $\leq 2$ ("low-confidence dataset") | 490 | 129 |
| Number of genes with multiple species carrying pCFM | 15 | 12 |
| Number of genes with multiple species carrying pCFM that passed all the quality filters ("high-confidence dataset") | 5 | 6 |

**Table 2.**
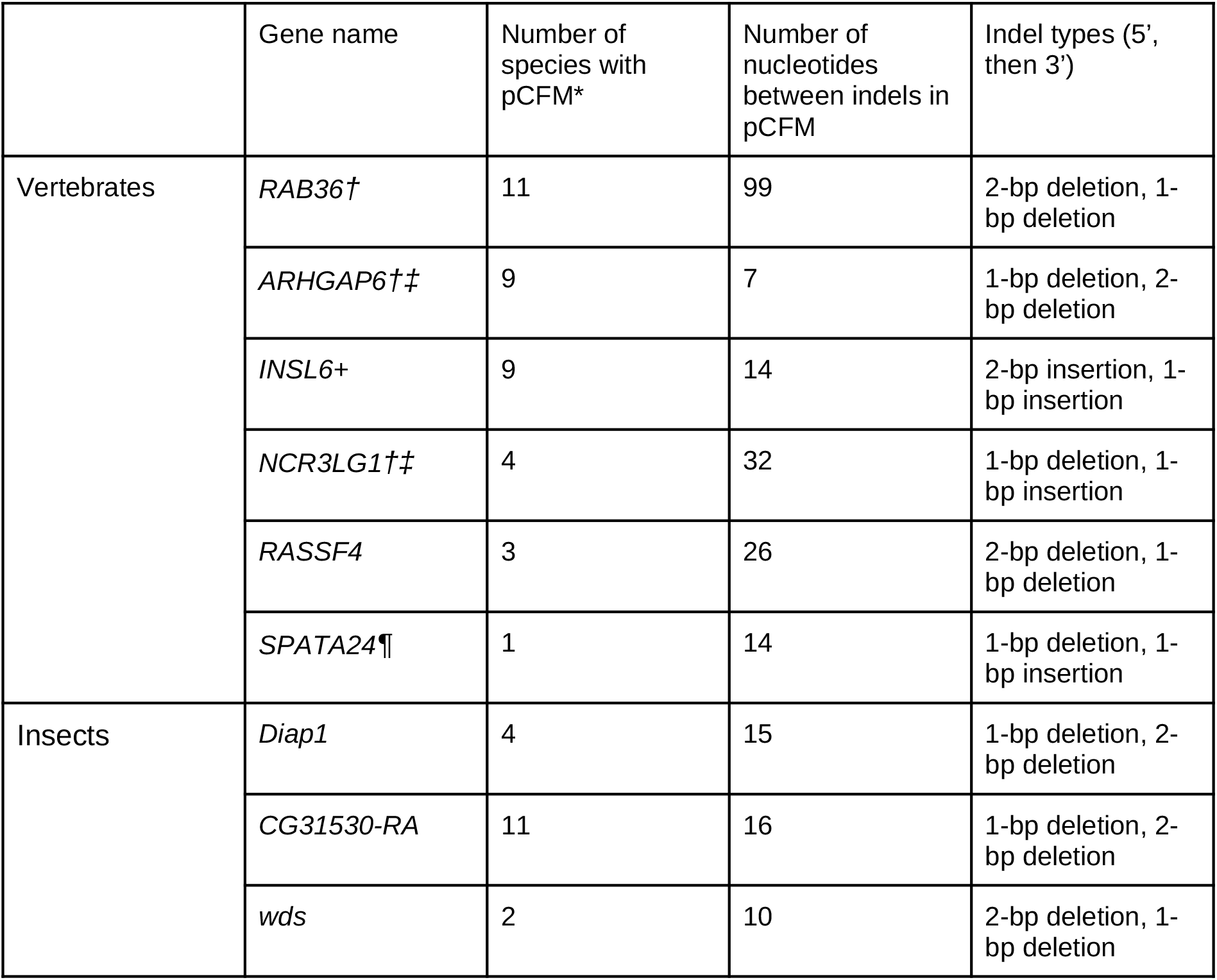
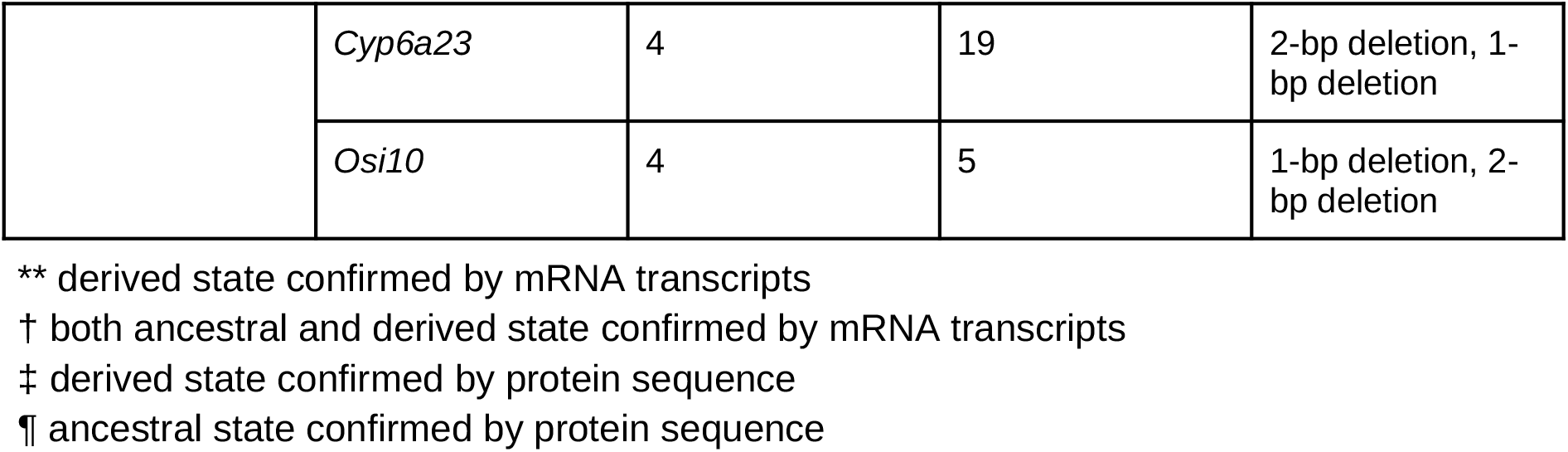
The 11 pCFM-carrying genes in the high-confidence dataset. The number of species carrying the pCFM (*) includes the species that were parts of the post-frameshift clades, but that were excluded at the filtering step (see the “Inference of pCFMs” sectio in Materials & Methods). For *SPATA24*, a closely related pCFM-carrying species (*Cavia aperea*) not present in the original alignment was found in ENSEMBL; therefore, this gene was included in the final list. See Supplementary Table 1 for lists of pCFM-carrying species for each pCFM.

**Figure 1.**
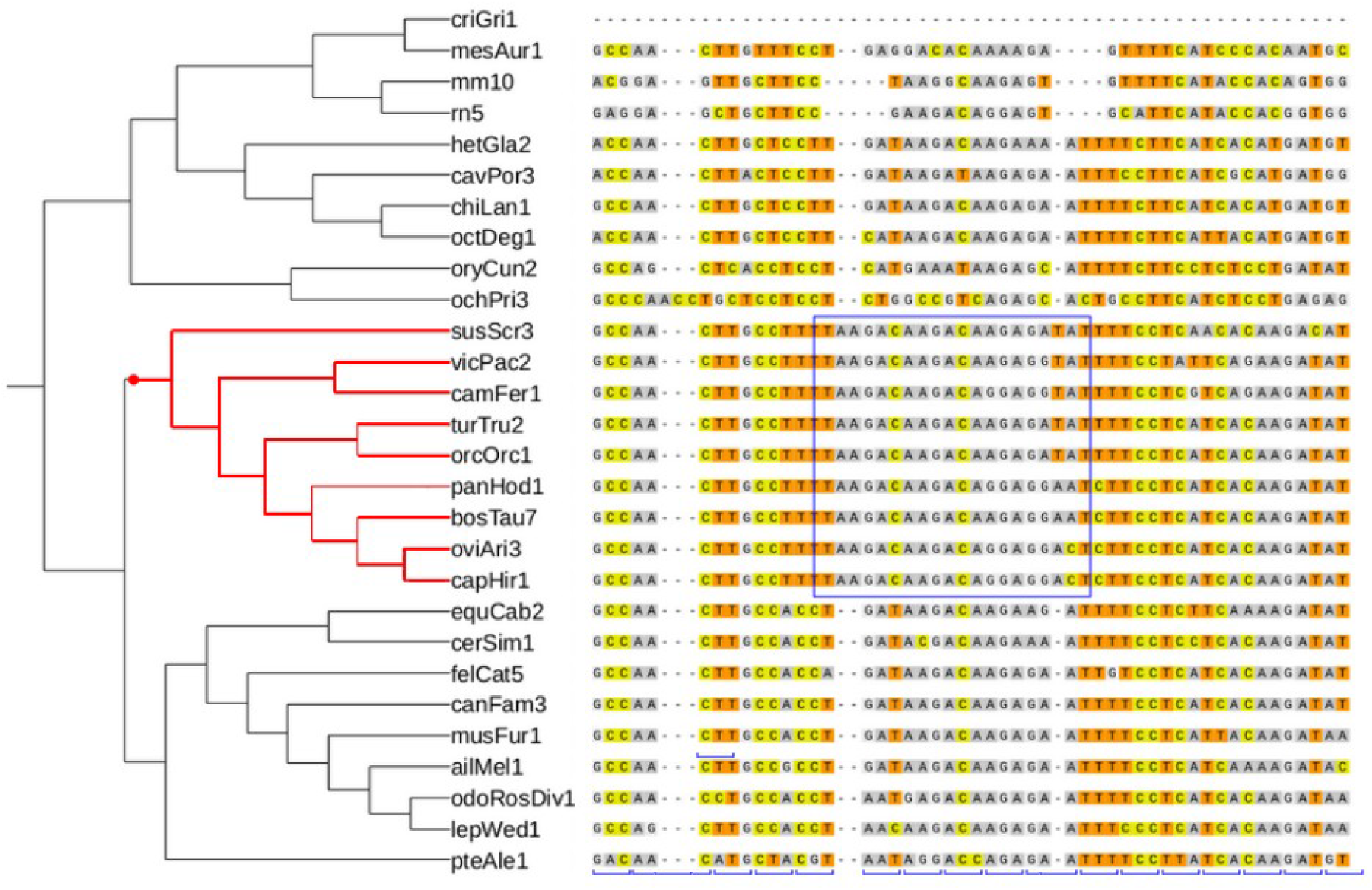
An example of a pCFM: a fragment of alignment of the *INSL6* gene, one of the 11 identified genes carrying high-confidence pCFM. At the left, a cladogram of the corresponding species is shown, with the pCFM-carrying clade marked in red, and a subset of the species devoid of the pCFM, in black (some of the non-pCFM-carrying species are not shown). The reconstructed phylogenetic position of the pCFM is marked with a red dot. The region with the shifted reading frame is marked by a blue rectangle. The original reading frame is indicated at the bottom of the alignment by blue square brackets.

A previous work (Hu & Ng 2012) has identified a pCFM between human and dog in the FLJ43860 gene. However, we were unable to confirm this finding, as the corresponding amino acid-coding sequence is not detected by tblastn in the reference human genome, and this gene is currently annotated as a pseudogene in HGNC gene symbol report (Povey et al. 2001).

### pCFMs are overrepresented near gene ends

Uncompensated indels with lengths not in multiple of 3 occur more frequently near ends of coding sequence, probably because the deleterious effect of a frameshift is weaker at these positions (Hu & Ng 2012; MacArthur et al. 2012). The pCFMs could follow the same pattern; alternatively, they could be distributed along the sequence uniformly as they do not cause any disruption of the protein-coding potential of the sequence downstream of them.

We find that pCFMs from the low-confidence dataset are biased towards gene ends (Figure 2). To quantify this, we compared the distribution of pCFM positions along the CDS with two other distributions: a uniform distribution, and that of uncompensated indels in human polymorphism data obtained from (Ng et al. 2008) (Figure 2). Specifically, we obtained the relative position of the pCFM along the CDS sequence as 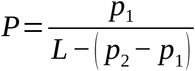, where *p*_1_ , *p*_2_^▫^are the positions of the 5’ and the 3’ indels and *L* is the length of the gene; *P*=0 when the 5’ indel is positioned at the 5’-most end of the CDS, and *P*=1 if the 3’ indel is positioned at the 3’-most end of the CDS. The distribution of *P*within a coding sequence significantly deviates from the uniform distribution (Kolmogorov-Smirnov test, p-value = 0.001), but does not differ from the distribution of indels in human polymorphism (Kolmogorov-Smirnov test, p-value = 0.12).

**Figure 2.**
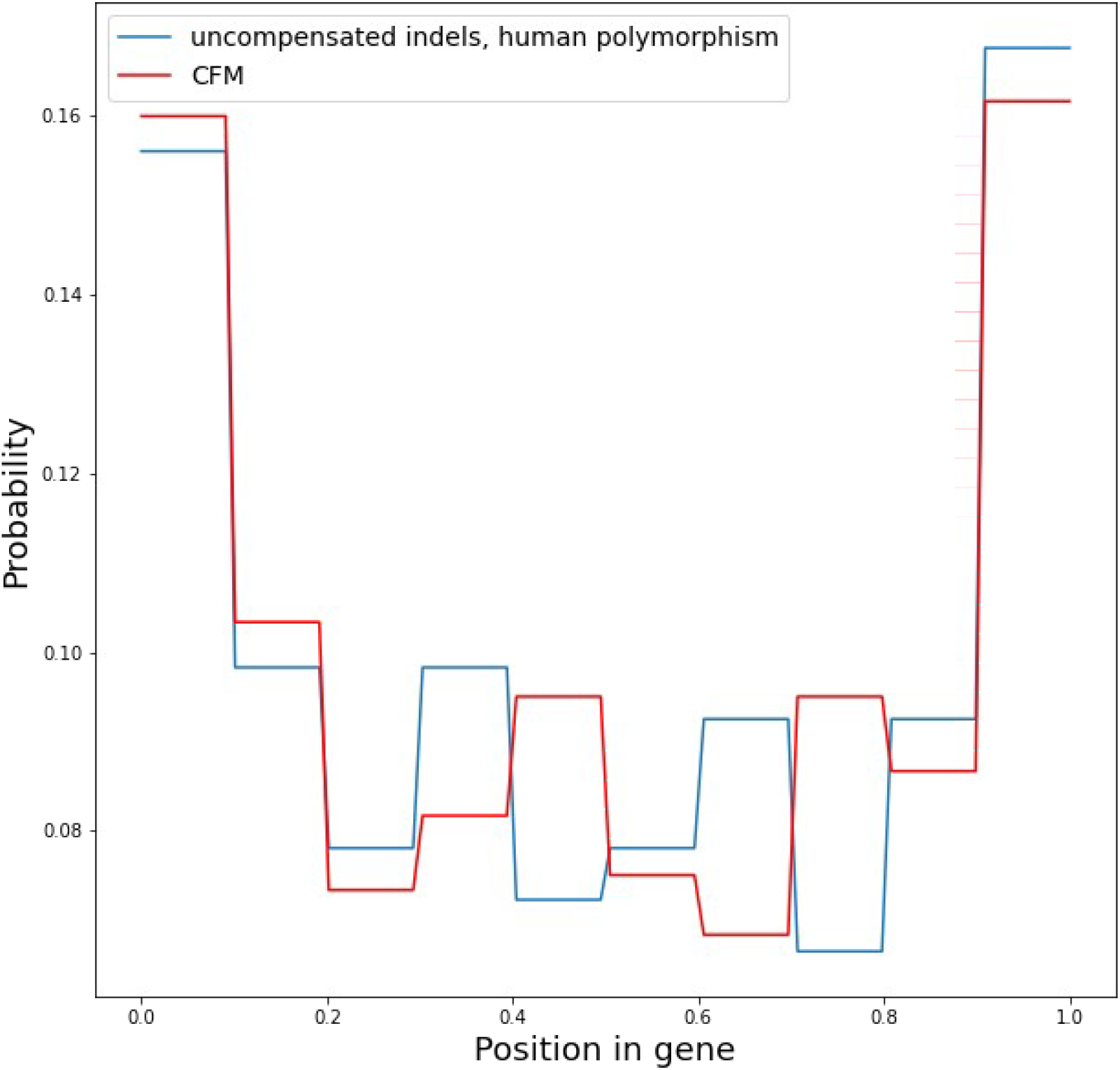
The distribution of uncompensated indels in human polymorphism data (blue line) and pCFMs (red line) along the gene coordinates. The horizontal axis indicates the position along the length of the coding sequence, with 0.0 corresponding to its start, and 1.0, to its end.

### Genes with compensatory frameshifts are under relaxed negative selection

We hypothesized that compensatory frameshifts are more likely to fix in genes that are under relaxed negative selection. To test this, we estimated the ratio of nonsynonymous to synonymous substitution rates ω for the pCFM-carrying genes, and compared it to the same value calculated for non-pCFM-carrying genes (Figure 3).

**Figure 3.**
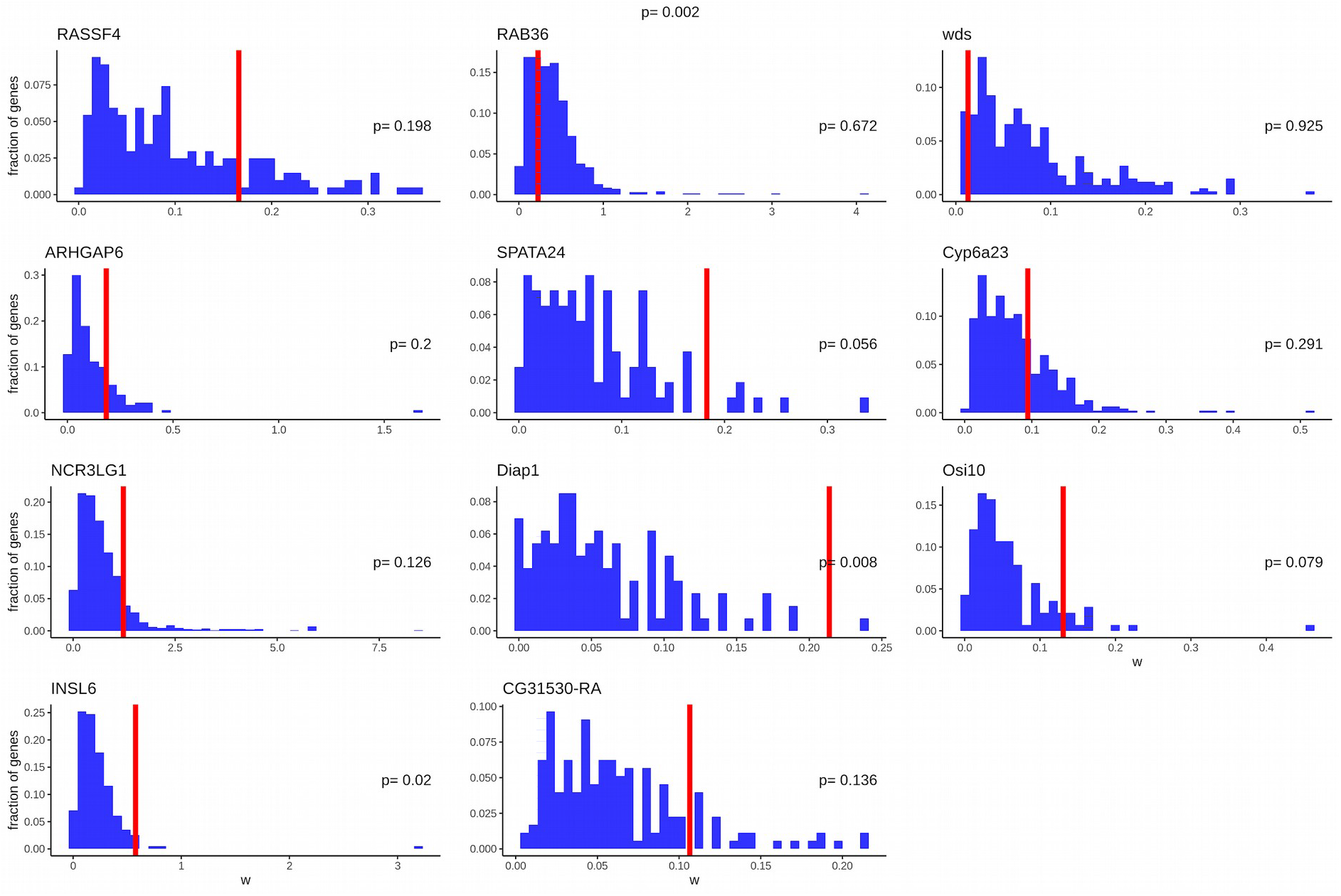
Negative selection at pCFM-carrying genes. Each panel corresponds to a gene with a pair of compensatory frameshifts. The red line indicates the observed value of the tree-wide ω of the gene of interest. The blue histograms show the distribution of ω in non-pCFM-carrying genes. The p-values are obtained as the proportion of the distribution to the right of the red line. Because ω were inferred using only those species that passed all quality filters, the numbers of species used to calculate ω differed among genes, which could affect ω estimates; therefore, for each panel, only those genes that had the same number of species were used for the reference distribution (see “Statistical approaches” section in Materials & Methods). The p-value of Kolmogorov-Smirnov test for uniform distribution of p-values is shown at the top.

The values of ω were above the 95th percentile of the distribution, indicating relaxed negative selection for two of the 11 genes from high-confidence dataset, *INSL6* and *Diap1*. They were also somewhat elevated for all genes of the high-confidence dataset combined (Kolmogorov-Smirnov test for uniform distribution of p-values, p-value = 0.002), although not for the low-confidence dataset (p-value = 0.53).

We also hypothesised that the mode of selection changes when the pCFM is fixed. To test this, for each gene, we fit two ω values: one for the clade of the phylogenetic tree descendant from the pCFM-carrying branch, and another for the remaining branches. To test whether this model describes our data better than the model with a single ω, we used the likelihood-ratio test. The two-ω model fitted the data significantly better than the null single-ω model for one of the genes (CG31530-RA in insects); for the remaining genes, no significant difference was found (Table 3).

**Table 3.** Selection in pCFM-carrying genes under different evolutionary models. ω, the ratio of nonsynonymous to synonymous substitution rates. See text for model details. *p<0.05 after the Bonferroni correction (p-values themselves are presented without a correction).

|  | One-parameter | Two-parameter model | Comparison of the |
| --- | --- | --- | --- |

Table 3. Selection in pCFM-carrying genes under different evolutionary models. $\omega$ , the ratio of nonsynonymous to synonymous substitution rates. See text for model details.
| Protein | model |  |  | model fits |  |
| --- | --- | --- | --- | --- | --- |
| | $\omega$ for the one-parameter model | $\omega$ for the non-pCFM-carrying branches | $\omega$ for the pCFM-carrying clade | LRT statistics value | LRT p-value |
| RASSF4 | 0.166 | 0.163 | 0.254 | 3.78 | 0.05 |
| NCR3LG1 | 1.23 | 1.71 | 0.867 | 1.76 | 0.183 |
| RAB36 | 0.229 | 0.24 | 0.211 | 0.22 | 0.636 |
| SPATA24 | 0.183 | 0.181 | 0.284 | 0.734 | 0.391 |
| INSL6 | 0.578 | 0.605 | 0.438 | 2.04 | 0.153 |
| ARHGAP6 | 0.184 | 0.184 | 0.187 | 0.01 | 0.912 |
| Diap1 | 0.214 | 0.213 | 0.235 | 0.207 | 0.649 |
| CG31530-RA | 0.107 | 0.101 | 0.155 | 11.013 | 0.001* |
| wds | 0.013 | 0.013 | 0.015 | 0.128 | 0.721 |
| Cyp6a23 | 0.094 | 0.094 | 2.129 | 0 | 1 |
| Osi10 | 0.131 | 0.128 | 0.177 | 3.45 | 0.063 |

### Protein regions spanned by pCFMs show no evidence for reduced conservation

To study the positions of pCFMs within the protein-coding sequence, we hypothesized that they tend to fix in those gene segments that are generally less conserved. To test this hypothesis, we compared the amino acid-level conservation of the region spanned by a pCFM to that of other CDS segments of the same gene, using a sliding window of the same length as the region spanned by a pCFM to obtain the null distribution (Figure 4). We used Shannon’s entropy as a measure of conservation of an amino acid site. We see no obvious decrease in conservation of the region spanned by a pCFM for the low-confidence dataset (Kolmogorov-Smirnov test for uniform distribution of p-values, p-value = 0.194) or the high-confidence dataset (Kolmogorov-Smirnov test for uniform distribution of p-values, p-value = 0.556).

**Figure 4.**
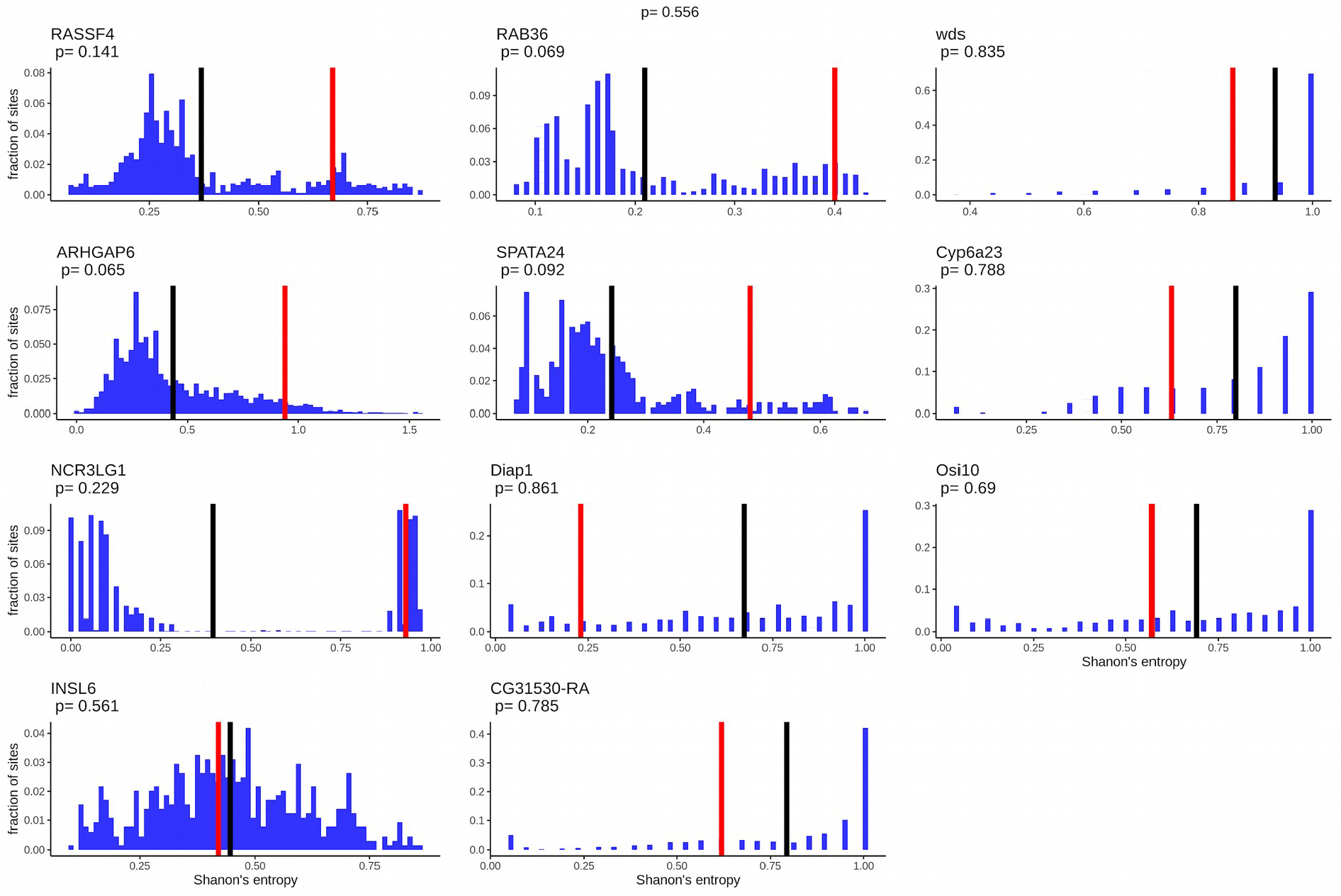
Conservation of the region spanned by a pCFM. Each panel corresponds to a gene with a pair of compensatory frameshifts. The red line indicated the mean Shannon’s entropy of the region spanned by a pCFM. The blue histograms show the distribution of entropy at all positions of the same protein, calculated as the mean entropy in a sliding window of length equal to the length of the region spanned by a pCFM, with the mean across all sliding windows indicated with the black line. The p-value of Kolmogorov-Smirnov test for uniform distribution of p-values is shown at the top.

### The effect of pCFMs on the encoded amino acid sequence

Due to the structure of the genetic code and the characteristics of the structured amino acid sequences, frameshifting mutations generally preserve the physico-chemical properties of the encoded protein (Wang et al. 2016; Bartonek et al. 2020). We asked if the pCFMs additionally tend to occur at those positions where this preservation is more precise.

Specifically, we hypothesized that pCFMs tend to fix at positions where they don’t substantially affect the structure of the encoded protein according to two similarity metrics. To test this hypothesis, for each of the 11 pCFM-carrying genes from the high-confidence dataset, we calculated the similarity between their amino acid sequences before and after the pCFM (see “Similarity metrics” subsection in Materials & Methods). We used ancestral state reconstructions (see “Ancestral states reconstruction” subsection in Materials & Methods) to infer the sequence in which the pCFM has fixed. We asked if the sequence resulting from a pCFM was unexpectedly similar to its ancestral version prior to the pCFM. To calculate the expected distribution of the similarity values, we introduced *in silico* pCFMs in different genes sampled randomly from the genome, preserving the indel types (insertion/ deletion), lengths (1 or 2) and the number of nucleotides between them.

We used two measures of sequence similarity between amino acid sequences: the Miyata distance and differences in hydropathy index. However, we present here only the result for Miyata distance for brevity (Figure 5); all the results on hydropathy index difference can be found in Supplementary materials (Supplementary figures 3-5). For individual genes, the observed effect of pCFM on the amino acid sequence matched the expectations, with none of the genes having significantly lower-than-expected differences between the ancestral and derived sequences after the Bonferroni correction. Overall the effect of pCFMs across all genes is also absent (Kolmogorov-Smirnov test for uniform distribution of p-values, p-value = 0.093 and 0.164).

**Figure 5.**
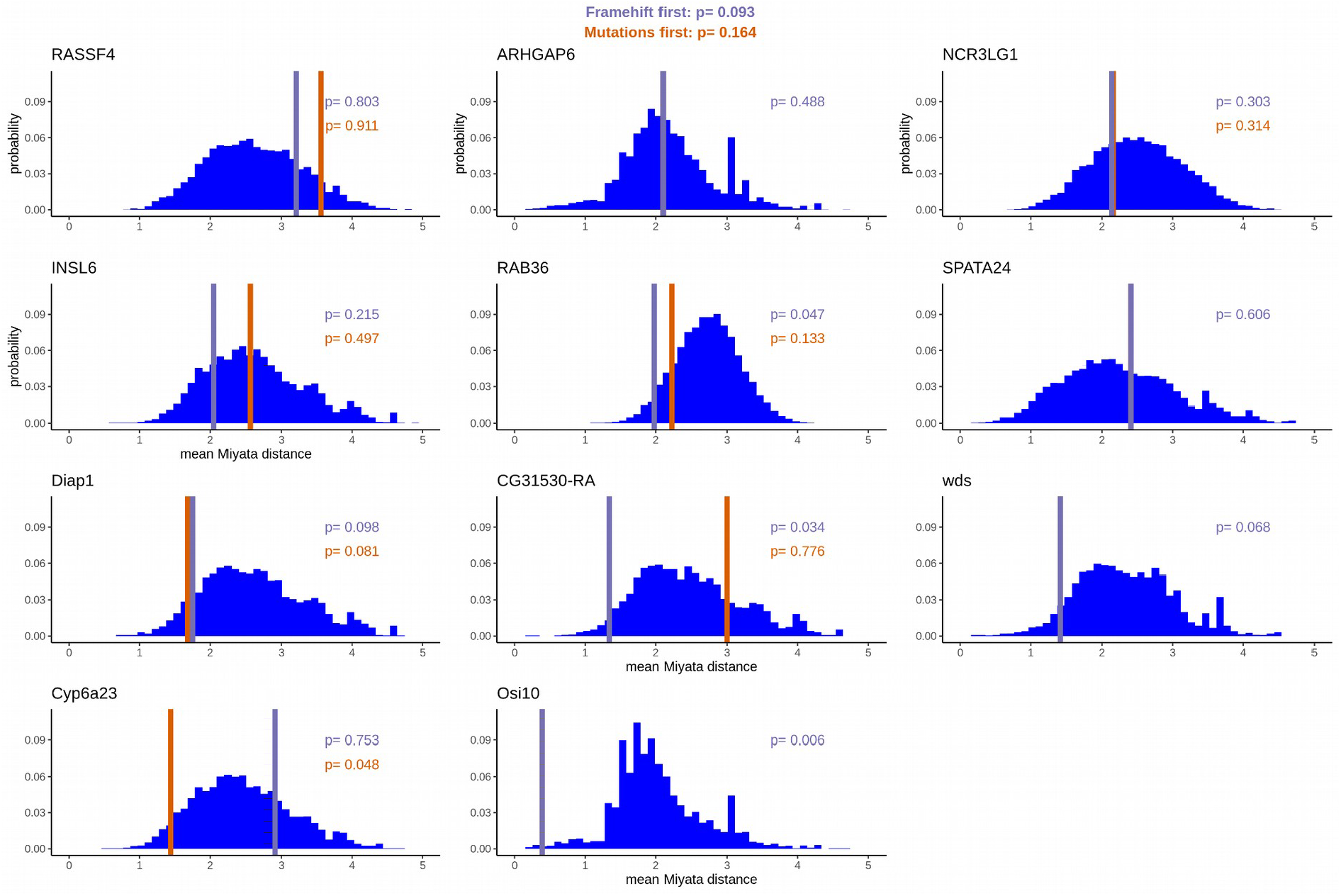
The effect of pCFMs on the physico-chemical properties of the encoded amino acid sequence as measured by the mean Miyata distance. The vertical lines represent the Miyata distance between the reconstructed ancestral sequence of the gene immediately prior to the pCFM, and the reconstructed (or observed) derived state of this sequence immediately after the pCFM. The purple and orange lines differ in treatment of single-nucleotide substitutions that have happened on the same phylogenetic branch as the pCFM: such substitutions are assumed to have occurred either after (purple line) or before (orange line) the pCFM; if no such substitutions were present, only the purple line is shown. The blue bars represent the distribution of Miyata distances between each of the 10,000 protein-coding sequences randomly drawn from the genome, and a version of this sequence with a pCFM spaced identically to that in the considered gene. The numbers in each panel correspond to the p-values obtained as the percentile of the distribution, with low p-values corresponding to lower-than-expected Miyata distances. The p-values of Kolmogorov-Smirnov test for uniform distribution of p-values are shown at the top.

### Interaction between pCFMs and single nucleotide substitutions

Conceivably, changes in physico-chemical properties of a protein segment caused by a pCFM could be partially compensated by subsequent single-nucleotide substitutions in the region spanned by them, if such substitutions make the protein more similar to the pre-pCFM state. Additionally, a pCFM could preferentially occur in regions where preceding amino acid substitutions made its effect less pronounced. To study this, we analyzed the interactions between pCFMs and substitutions that happened on the same phylogenetic branch in their effect on the characteristics of the encoded amino acid sequence.

In Figure 6, we measure the amino acid similarity between the reconstructed ancestral variant of the protein (A), the variant carrying pCFM (A_fr_), the variant carrying the single-nucleotide substitutions that occurred at the same branch as pCFM (A_mut_), and the variant carrying both pCFM and the substitutions (E). The observed patterns of pairwise similarity between these variants differed between proteins.

**Figure 6.**
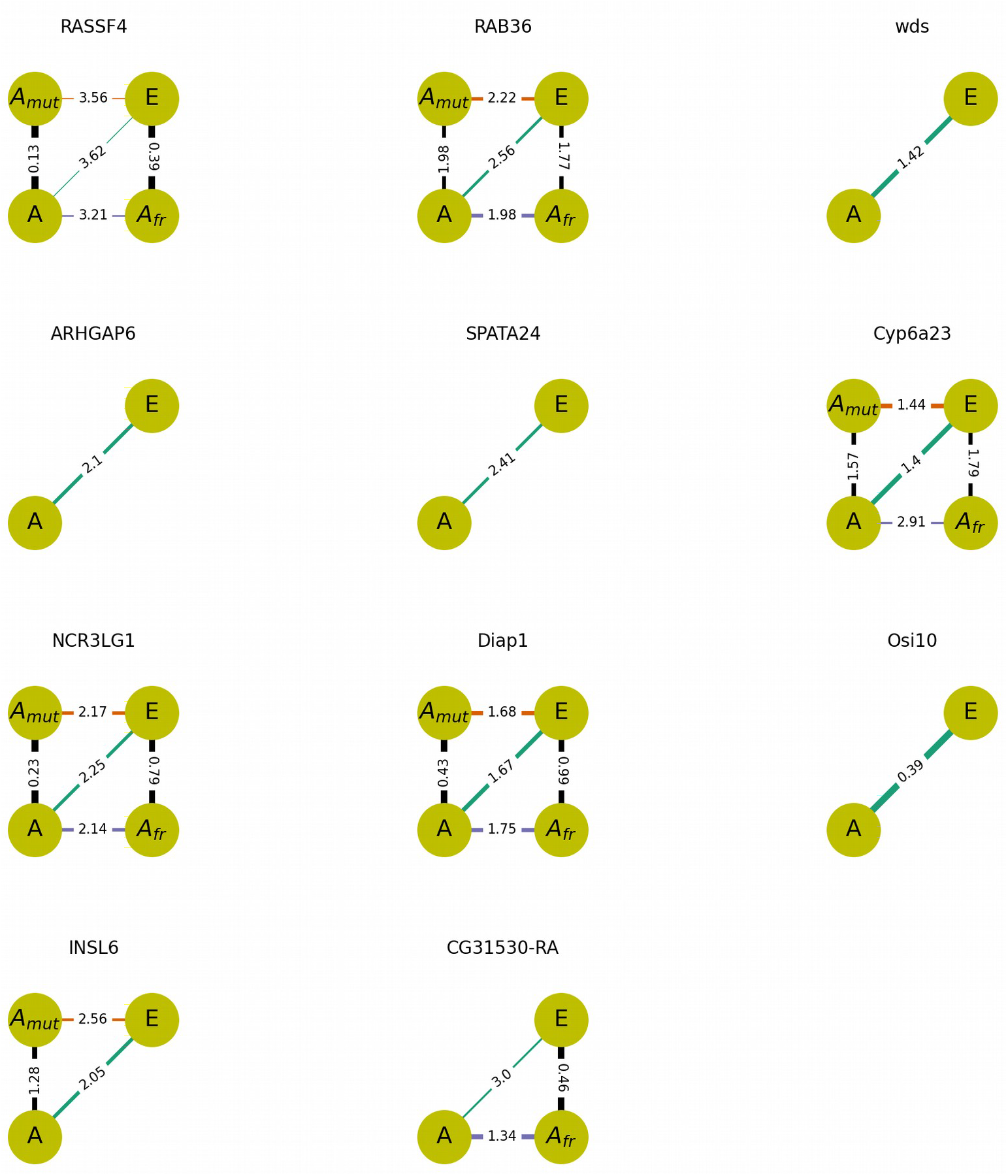
Effects of a pCFM and amino acid substitutions occurring on the same phylogenetic branch as pCFM on the physico-chemical properties of the encoded protein, according to the Miyata difference between the encoded amino acid sequences. The four circles in each panel represent the four sequences: without pCFM or substitutions (ancestral state, A); with substitutions but without pCFM (A_mut_); with pCFM but without substitutions (A_fr_); with both pCFM and substitutions (derived state, E). The numbers on the lines connecting these states represent the distances between corresponding sequences; smaller distances are depicted with bolder lines. Omitted intermediate states (A_mut_ or A_fr_) correspond to cases when the substitutions are synonymous in the context of pCFM and nonsynonymous in the context of the ancestral state, or vice versa. Both intermediate states are omitted if no amino acid substitutions happened on the considered branch.

Importantly, we were unable to infer the order in which pCFM and substitutions occurred on the same branch. Therefore, we had to consider two possibilities: that pCFM occurred prior to the substitutions, and that pCFM occurred after the substitutions. We put forth two hypotheses with regard to the similarity patterns: (i) that the substitutions occurred after the pCFM, and compensated for its effect; (ii) that the substitutions occurred prior to the pCFM, and made it permissible, so that the effect of the pCFM was weaker on the mutated background.

Under (i), we expect the (A, E) distance (green in Figs. 7-8) to be lower than the (A, A_fr_) distance (purple). This was the case for substitutions in *RASSF4* and *NCR3LG1* by hydropathy measurements, in *Diap1*, by Miyata distance measurements, and in *Cyp6a23*, by both measurements.

**Figure 7.**
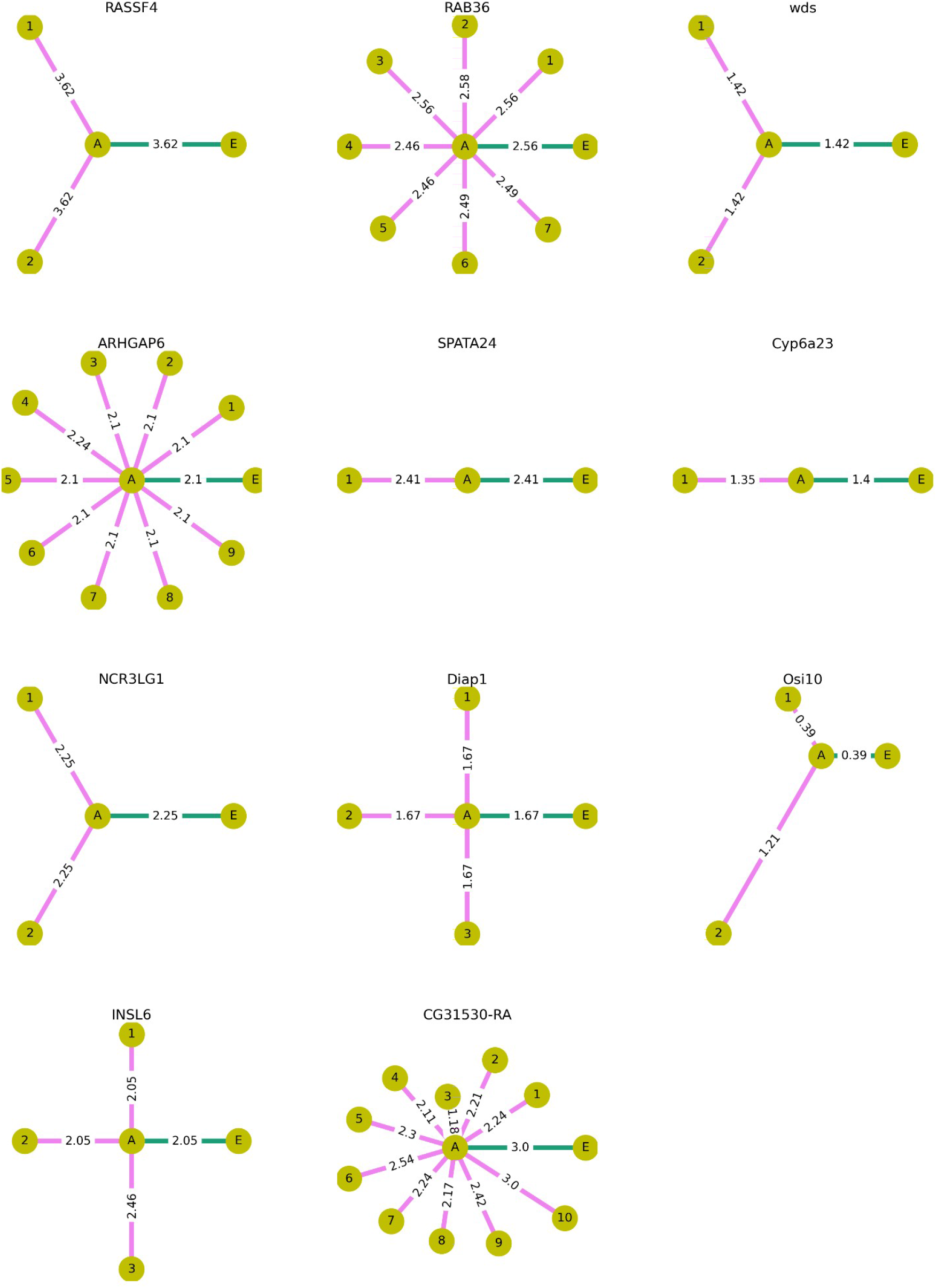
Effects of amino acid substitutions occurring on the phylogenetic branches descendant to pCFM on the physico-chemical properties of the encoded protein, according to the Miyata difference between the encoded amino acid sequences. In each panel, the circles correspond to the different states of a sequence: without pCFM (A); with pCFM (E, always to the right of A); and with both pCFM and mutations on the descendant branches (terminal leaves states, denoted by numbers). The lengths of the lines and the numbers on top of them represent distances between the corresponding states; many of the distances are identical, reflective of the substitutions common to the corresponding species.

Under (ii), we expect the (A_mut_, E) distance (orange) to be lower than the (A, A_fr_) distance (purple). This was the case for substitutions in *RASSF4, INSL6* and *NCR3LG1* by hydropathy measurements, and in *Diap1* and *Cyp6a23* by Miyata distance measurements.

Nevertheless, we observe no general tendency for the substitutions to be compensatory or permissive.

Next, we hypothesized that the deleterious effect of pCFM can be compensated by substitutions occurring at subsequent branches in the phylogeny (Figure 7). If this hypothesis holds true, we expect the distance between A and terminal descendant branch (magenta in Figure 7) to be smaller than the (A, E) distance (green). This was the case in *RAB36, CG31530-RA* and *Cyp6a23* by Miyata distance measurements and in *ARHGAP6, RAB36, CG31530-RA* and *Osi10* by hydropathy measurements (Supplementary Figure 5). Overall, however, we see no tendency for post-pCFM substitutions to be compensatory: while 14 of them decrease the distance to the ancestral sequence by Miyata distance (13 by hydropathy measurements), 4 increase it (7 by hydropathy measurements), and the 14 cases of decrease in Miyata distance are highly correlated because of the tree structure.

## Discussion

In a recent paper, Bartonek et al. (Bartonek et al. 2020) indicated that frameshifts tend to preserve the physico-chemical properties of the encoded amino acid sequence, and asked: “can evidence be found that frameshifting has indeed played a relevant role during evolution of real proteins?”. Here, we address this question, and show that this is indeed the case. We describe 129 instances in the recent protein evolution of insects, and 490 cases in vertebrates, where a pCFM has fixed in the course of evolution of the lineages that gave rise to extant species. By using stringent filtering criteria, we focus on the 6 insect and 5 vertebrate genes for which this evidence is the most robust, and study how they affect the encoded amino acid sequence. Importantly, we focus on pCFMs that happened rapidly one after the other (so that they were observed on the same segment of the phylogenetic tree); thus, we ignore any possible instances of a longer time lag between the two events, including possible cases of resurrection of established pseudogenes (Esfeld et al. 2018).

Among our pCFMs consisting of two same-type events (insertion-insertion or deletion-deletion), the deletion-deletion pairs are more frequent (Table 2), and this pattern is also observed in the larger unfiltered dataset. This is consistent with the fact that, at least in *Drosophila*, deletion mutations are more frequent than insertion mutations (mutational deletion bias). In functional regions, this bias is compensated by a higher probability of insertions to spread and be fixed by selection (fixational insertion bias) (Leushkin et al.2013). As a result, among fixed in-frame exonic mutations, insertions and deletions are equally frequent (Fig. 2 in (Leushkin et al. 2013)). The fact that the deletion bias is retained for our fixed pCFMs suggests that unlike in-frame indels, selection against out-of-frame insertions and deletions is similar, so that the excess of deletions among fixed mutations recapitulates the biases at origin of these mutations.

### pCFMs and sequence conservation

The 11 high-confidence genes with fixed pCFMs are generally weakly constrained, as suggested by their lower ω values. This is in line with the expected radical effect of frameshifting indels on the encoded protein, and with the majority of them being deleterious. This trend is not reproduced in the low-confidence dataset, potentially because it might be contaminated by false positives.

The deficit of pCFMs at more constrained proteins could arise for two non-mutually exclusive reasons. It could be due to selection against the intermediate variant carrying a single frameshifting mutation (which would have to be present at the population at some point unless both mutations occurred simultaneously in the same individual); it could also arise from selection against the ultimate pCFM variant.

The finding that pCFMs do not preferentially fix in the least conserved regions of a protein favors the first explanation. Indeed, it suggests that the overall constraint of the protein imposes stronger limitations on the pCFM than the constraint of the region spanned by the pCFM: pCFM may fix in any region if the protein is sufficiently unimportant. This would be the case if the limiting step was survival of some individuals with uncompensated frameshift for some number of generations. By contrast, if the pCFM positions were mainly determined by the effect of selection at the pCFM stage, we would expect pCFMs to preferentially happen at regions of low conservation, as such selection would discriminate between the protein versions with ancestral and derived amino acid sequences encoded by the region spanned by pCFM.

The observation that pCFM distribution along the gene follows that of uncompensated indels also supports the hypothesis that the intermediate state is limiting. Indeed, as the overall selective constraint is generally rather uniform along the CDS length (Shabalina et al. 2004), the same uniformity is expected for pCFMs if the intermediate uncompensated state is not selected against. By contrast, we expect the distribution of pCFMs to follow the distribution of uncompensated indels if this pattern is produced by selection against the state carrying a single frameshifting mutation.

Together, these findings suggest that selection has enough time to act against the single-indel mutant, probably while it segregates at low frequencies within the population.

The fact that pCFMs preferentially fix in genes under relaxed negative selection does not mean that the pCFMs are functionally unimportant. In one of the high-confidence genes (CG31530-RA), we find that negative selection has relaxed at the same time as the pCFM was fixed, suggesting that the pCFM could have perturbed its function. This gene encodes a transmembrane protein involved in transmembrane transport (UniProt Consortium 2008; https://www.uniprot.org/uniprot/D2NUL5). *Drosophila melanogaster* and 10 other species of flies have the version of this gene with indels.

### Expected frequency of pCFMs

In a functional gene, a single frameshifting mutation may cause a loss in fitness, which can then be compensated by the second frameshifting mutation. There is extensive theoretical literature aiming to estimate the possibility and rates of compensatory evolution under various ranges of population genetics parameters. To our knowledge, the first such model was proposed by Kimura (Kimura 1985); using diffusion approximation, he showed that the rate of compensatory evolution at tightly linked loci is non-negligible over a wide range of *N*_e_*s*, where *N*_e_ is the effective population size and *s* is the coefficient of selection disfavoring the intermediate variant. This approach was further developed by Stephan (Stephan 1996) who focused on RNA evolution and expanded the model assumptions to different selection coefficients against different intermediate mutants.

In subsequent work, two distinct mechanisms for such valley crossing were considered. The first one is sequential fixation of mutations: the first (single or multiple) mutations are deleterious and fixed by drift, and the last mutation is advantageous against their background and is fixed by selection. Under the second mechanism, the intermediate deleterious variant is never fixed; instead, the haplotypes carrying it remain at low frequencies, either due to genetic drift (so-called stochastic tunneling (Iwasa et al. 2004)) or deterministically under mutation-selection balance, before they are “rescued” by subsequent compensatory mutations. The range of parameters leading to each regime was explored by Weissman (Weissman et al. 2009) in asexual populations; for sexual populations, he has shown that low recombination rates may even facilitate valley-crossing (Weissman et al. 2010). One of the results of this study was that, quite intuitively, tunneling is relevant for larger *N*_e_, while sequential fixations are relevant for smaller *N*_e_. Carter and Wagner (Carter & Wagner 2002) also obtained similar results, invoking differences in *N*_e_ to explain the differences in compensatory evolution rates of enhancers in vertebrates compared to

#### Drosophila

The excess of pCFMs near gene ends, the independence of the pCFMs frequency of the conservation of the corresponding CDS segment, and the higher frequency of pCFMs in weakly selected genes together suggest that fixation of a pCFM proceeds through a polymorphic intermediate state. To test whether this assumption is consistent with data, we adopt a simplistic model to estimate the number of pCFMs we could expect to find in vertebrate and mammalian genome-scale datasets. We consider two tightly linked positions such that mutations are deleterious in each of them separately but are neutral in combination. The selection coefficient against both single mutants *s* is assumed to be the same and to equal that against a loss of function in a non-essential gene. This selection is assumed to be strong, such that individual frameshifting mutations are unlikely to fix in a population under realistic *N*_e_. Therefore, we only consider the rate of tunneling, assuming that the first frameshifting mutation is maintained by mutation-selection balance before the second, compensatory mutation occurs. These assumptions are close to those of Kimura’s (Kimura 1985).

Let *L* be an average gene length; *p*_1_, the rate of the first uncompensated mutation per nucleotide per generation; and *p*_*2*_, the rate of the second, compensatory mutation per nucleotide per generation. The probability of a frameshifting mutation at some position of this gene per generation *m*_1_ then equals

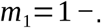

Similarly, the probability of a compensatory mutation at some position of this gene per generation *m*_2_ equals

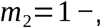

where *l* is the number of nucleotides surrounding the first mutation where the second mutation would be compensatory.

Under mutation-selection balance (Crow & Kimura 1970), the expected frequency of uncompensated frameshift in a population equals *m*_1_/*s*. The compensated genotypes will then originate by mutation of the deleterious alleles at per generation rate of *m*_1_ *m*_2_/ *s*.

If compensation is exact, so that the compensated genotype has the same fitness as the ancestral variant, the rate of fixation of such compensatory pairs equals the rate of the mutation giving rise to them: *P*_*fix*_=*m*_1_ *m*_2_ / *s* (Kimura & M. 1983). Over *T* generations, a total of *T m*_1_ *m*_2_ /*s* such pairs are expected to have occurred.

To estimate *p*_1_and *p*_2_, we consider four types of deleterious frameshifting mutations: insertion of lengths 1 or 2 and deletions of lengths 1 or 2. For each of these mutations, we consider the sets of mutations that can compensate for it (for example, an insertion of length 1 can only be compensated by an insertion of length 2 or a deletion of length 1 but not by a deletion of length 2), and sum over the probabilities of these mutations. The resulting *P*_*fix*_ terms are the sums over these pairs of deleterious and compensatory mutations.

We use the estimate for the *de novo* mutation rate *μ*=10^*-* 8^ for vertebrates from (Kong et al. 2012) and *μ*=4.9 ∗10^*-*9^ for *Drosophila* from (Assaf et al. 2017); the estimates for *p*_1_ for each type of frameshifting mutation from (Leushkin et al. 2013):0.020 *μ*,0.010 *μ*,0.042 *μ*and 0.015 *μ* for insertion of length 1, insertion of length 2, deletion of length 1 and deletion of length 2, respectively; and the estimate for the selection coefficient against a heterozygous loss of function in non-essential genes *s*=0.0015in *Drosophila (Langley et al. 1981)* and *s*=0.005for vertebrates (estimated for human in (Cassa et al. 2017)). We estimated *T* and *L*from our data. For each gene, *T* the mean length of a phylogenetic tree including only the species that have passed all filtering steps, without accounting for the terminal branches (since we only consider indels inherited by multiple species) and measured in the genome-wide number of synonymous substitutions. In our data, *T* =176536337 for vertebrates, and *T* =403272889 for insects. *L* is just the mean length of analysed genes, and equaled *L*=1897 for vertebrates and *L*=1204 for insects. We take *l* to equal 100 (the size of all the regions spanned by observed pCFMs is less than that, Table 2).

Using these estimates, we obtain *Tm* 1 *m* 2/ *s*=0.002for vertebrates and0.003for insects. Given the number of protein-coding genes we analyzed (21208 for vertebrates and 15283 for insects), we expect to find 52 ± 14 and 43 ± 13 genes with pCFMs in vertebrates and insects respectively (95 credible intervals are calculated using normal distribution approximation for binomial distribution with parameters *n*=21208, *p*=0.002 and *n*=15283, *p*=0.003).

The above values lie between the numbers of raw unfiltered indels (490 genes in vertebrates and 129 in insects) and filtered indels (6 and 5 respectively) that we obtain. This could indicate that our filtering steps are too stringent, perhaps excluding some of the *bona fide* pCFMs. It could also result from violations of our assumptions. In particular, the above model assumes that a fixed pCFM confers the same fitness as the wild type sequence; if it is selectively inferior, the expected numbers will be lower, and if it confers a selective advantage, these numbers will be higher.

### Conclusions

We describe 619 candidate pairs of compensatory frameshifting mutations in the past evolution of vertebrate and insect species, including 11 high-confidence cases that passed all our stringent filtering steps. pCFMs tend to fix in genes under relaxed negative selection, but not in particularly poorly conserved segments, suggesting that the rate of their fixation is mainly limited by selection against individual uncompensated frameshifting mutations. In particular, this implies that pCFMs occur as a pair of distinct mutational events separated by some period of time, rather than a single mutational event of two simultaneous indels. This favors the model of origin of pCFMs under which a single deleterious frameshifting mutation segregates at a low frequency within a population until it is compensated by another indel mutation arising in the same haplotype. Overall, these results put forward pCFMs as a previously undescribed source of novelty in protein evolution.

## Materials & Methods

### Sequence alignment

As a starting point, we used the 100-way vertebrate and 124-way insect (diptera) MULTIZ alignments obtained from the UCSC genome browser (Rosenbloom et al. 2015). These alignments are comprised of non-reference genomes being aligned to a reference (correspondingly, human or *Drosophila melanogaster)*. This means that all the sites where the reference species has a deletion are absent, making these alignments unfit for our purposes. Therefore, we realigned the protein-coding sequences of genes from these species as follows. We extracted the exonic sequences from genomes using the annotation from the corresponding MULTIZ alignments, concatenated these exons into genes, and aligned the resulting genes using mafft v7 (Katoh et al. 2002). The following parameters were used both for vertebrate and for insects: --maxiterate 1000 --globalpair –preservecase. To allow for detection of frameshifts, we performed a nucleotide-level, rather than a codon-level, alignment. In the insects data set, we removed the D_pseudoobscura_1 and A_gambiae_1 genomes as they were redundant (the alignment already contained the droPse3 and anoGam3 genomes from the same species). Because the gene annotation was based on the genome of the reference species, we excluded the overhanging 5’-and 3’-ends as well as long (>50 nucleotides) internal DNA segments in non-reference species that had no orthologous sequence in the reference. High-quality alignments were obtained for 21208 out of 21521 genes of vertebrates and 27341 out of 30482 genes in insects (insect genes were comprised of all NCBI RefSeq Gene CDS regions including non protein-coding ones; non protein-coding genes were excluded at subsequent filtering steps). The phylogenetic trees for vertebrates and insects obtained from whole-genome alignments, with branch lengths scaled in estimated numbers of nucleotide substitutions per site, were obtained from the UCSC genome browser.

### Inference of pCFMs

We designed an algorithm for detection of pCFMs using a multiple alignment and the corresponding phylogenetic tree.

In short, the algorithm uses a parsimony-based approach to detect insertions and deletions in the alignment. As the first step, it reconstructs past insertions and deletions that had occurred at specific phylogenetic branches, and are inherited by subsets of analyzed species. At the second step, to ensure usage of high quality and high confidence protein-coding sequences for pCFM inference, the nucleotide alignments for each gene were filtered on a sequence-by-sequence basis. We excluded the species that did not meet all of the following criteria: had the number of nucleotides divisible by 3; carried a start codon at the sequence start; carried a stop codon at the sequence end; carried no in-frame stop codons. At the third step, pCFMs are identified as pairs of indels of length not divisible by 3 in the remaining sequences; at this step, we also excluded sequences with more than 2 such indels, assuming that our reconstruction of pCFMs in such sequences would be less trustworthy. We performed quality filtering after indel classification, assuming that the added information from the presumably lower-quality sequences allows for more robust classification of the evolutionary history of indels. For insect dataset alignments for which fewer than four sequences were left after filtering were not considered further; these included all non-protein-coding genes.

For each detected pCFM, the algorithm outputs their type (insertions or deletions), positions, lengths, and the list of species they were detected in. A detailed description of the algorithm is provided in the Supplementary text 1 (see also Supplementary Figures 1 and 2). The algorithm is implemented as a Python script, available at https://github.com/Captain-Blackstone/Compensatory-frameshifts.

### Post-filtering of pCFMs

With all the pCFMs detected, we then focused on the most reliable ones, narrowing down the considered sample as follows. First, while our algorithm detects pCFMs independently of whether they occurred in the same or in different branches of the phylogenetic tree, in this work, we only consider those that happened on the same branch. Second, we only considered pCFMs comprised of short indels, with combined lengths of no more than 4 nucleotides (i.e., two insertions of lengths 1 and 2; two deletions of lengths 1 and 2; an insertion and a deletion both of lengths 1; or an insertion and a deletion both of lengths 2). Third, the remaining alignments were manually inspected, and only those with unambiguous pCFMs were retained. This resulted in 619 pCFMs, contained in 619 distinct genes.

To obtain the high-confidence dataset, we focused on those cases where a pCFM was inherited by more than one species in our dataset. This way, we could have excluded some *bona fide* species-specific pCFMs, but obtained additional support for the remaining ones, under the logic that coincident artefacts in different independently assembled genomes should be rare. We also excluded those alignments in which either the ancestral or the frameshifted sequence was not supported by running nBLAST (Altschul et al. 1990) against the genome of the corresponding species in the NCBI nr database.

This left us with 10 high-confidence pCFMs. For the vertebrate dataset, we then sought additional support from other sequencing projects not included in the UCSC genome browser. For this, for each pCFM-carrying gene, we used ENSEMBL (Cunningham et al. 2019) to find orthologs in species that were more closely related to the pCFM-carrying species than any other species in our alignment. This rescued one additional pCFM for the final dataset, for a total of 11 pairs.

### Statistical approaches

For each gene, we used PAML version 4.8 (Yang 1997) to estimate the probabilities of nonsynonymous changes relative to synonymous ones (ω values) under the substitution model of Goldman and Yang (1994) described in Yang et al., 1998. We used the M0 model (model=0 parameter) to estimate a single ω value for the entire tree, and a model with two ω parameters (model=2 parameter) to fit distinct values of ω for the pre-and post-pCFM branches. The remaining parameters were the same in both models: runmode=1, seqtype=1, NSsites=0.

For each gene, we estimated ω using as input only the subset of species that passed the filtering steps described in the “Inference of pCFMs” section. As a result, the subset of considered species differed between genes.

To ask whether the pCFM-carrying genes are characterized by increased or decreased ω values, we compiled a sample of non-pCFM-carrying genes as a control. For this, we used the same filtering steps as above, namely, retained for each alignment only those sequences that had the number of nucleotides divisible by 3; carried a start codon at the sequence start; carried a stop codon at the sequence end; and carried no in-frame stop codon. To account for differences in numbers of considered species which could bias ω estimates, for each pCFM-carrying gene, we used as a control a subset of genes that were matched by the number of retained species.

To combine the signal across comparisons for individual genes, we used the Colmogorov-Smirnov test with the null hypothesis of the uniform distribution of p-values.

### Ancestral state reconstructions for nucleotide sites

Ancestral states for individual nucleotide sites and phylogenetic positions of single-nucleotide substitutions were reconstructed independently of the inference of indels (see above) with the help of the maximum likelihood method implemented in MEGA6 (Tamura et al. 2013). For substitutions that had happened on the same phylogenetic branches as the pCFM, which one came first could not be established unambiguously. In these cases, we assumed that all substitutions occurred simultaneously; that both frameshifting indels in the pCFM occurred simultaneously; and considered two scenarios, assuming that the substitutions occurred either before or after the pCFM. For these branches, we identified the ancestral state as “A”, the intermediate state in the substitution-first scenario as “Amut”, the intermediate state in the frameshift-first scenario as “A_fr_”, and the derived state as “E”.

### Similarity metrics

To study the effects of indels and single-nucleotide substitutions on the encoded amino acid sequence, we compared different states of these sequences using two similarity metrics. First, we measured the mean Miyata distance (Miyata et al. 1979) between the aligned amino acids in the pairwise alignment. This metric is also close to 0 for very similar sequences; for very distant sequences, it is close to 5.13, which is the maximum Miyata distance (namely, the distance between glycine and tryptophan). We considered gaps to have the maximum possible distance (5.13) to any amino acid. Second, we measured the difference between the mean hydropathy indexes of the two sequences (Kyte & Doolittle 1982). This difference is close to 0 for very similar sequences and close to 9 for very distant ones.

## Data availability

All processed data (after realignment and clipping) is available at FigShare (https://figshare.com/articles/dataset/Re-aligned_data_from_UCSC_100-way_vertebrate_and_124-way_insect_exome_alignments_/13272974). The python script used to infer pCFMs is available at the gitHub repository (https://github.com/Captain-Blackstone/Compensatory-frameshifts).

## Acknowledgements

The authors are grateful to Alexey Kondrashov for discussions of the model for the expected frequency of compensatory frameshifting mutations; to Sergey Lysenkov (Moskow State University) for advice on statistics; and to members of Georgii Bazykin’s and Alexey Kondrashov’s labs for valuable discussions. Computations were performed using the Makarich HPC cluster of the Faculty of Bioengineering and Bioinformatics, Lomonosov Moscow State University.

