## Supplementary Materials for "Pairs of mutually compensatory frameshifting mutations contribute to protein evolution in vertebrates and insects"

#### Supplementary Text 1: algorithm for detection of pCFMs

Here we present a complete description of the algorithm we used to infer compensatory frameshifting mutations from a nucleotide multiple alignment of a set of orthologous protein-coding sequences. The main steps are reflected in Supplementary figures 1 and 2.

1. All unique “holes” in the alignment are determined and the list of them is formed. The hole is defined as a row of successive gaps, which may further be classified as an insertion or a deletion. If multiple species have a row of successive gaps **of the same length** and **in the same position** they share a hole.

2. While the list of holes is not empty, paragraphs 2.1-2.4 are executed (holes are classified into insertions or deletions).

    2.1. The shortest hole is chosen, further referred to as the target hole; if multiple holes of the same minimum length are available, a random one of them is chosen.

    2.2. Two lists of species are formed. The first one includes species sharing the target hole. The second one includes species not sharing the target hole. Only species with non-gap symbols in all the positions of the hole are added to this group. *Note that a species carrying a longer hole in the same region is not considered to share the target hole.*

    2.3. For the two lists of species formed in paragraph 2.2, it is determined which of them form a clade (one of them, both or neither of them).

        2.3.1. If only the first group (with the target hole) forms a clade, the target hole is classified as a deletion in the species sharing it.

        2.3.2. If only the second group (without the target hole) forms a clade, the target hole is classified as an insertion in the species sharing it.

            2.3.2.a. If the target hole is classified as an insertion and there are other holes in the alignment overlapping with the target hole, the overlapping positions are removed from these holes. *This step assumes that all gaps in the species without an insertion at positions of this insertion are “insertion-induced” - they are there only because of the presence of species with insertion in the alignment. However, it may be not true, if a deletion appeared* ***after*** *the insertion and some of the inserted nucleotides got deleted as a result. In this case gaps may represent their own deletion instead of being insertion-induced. As far as we are concerned there is no way to know if gaps on the overlapping positions are insertion-induced or not. Therefore, such gaps were still considered insertion-induced, but were recorded as “uncertain cases”. Unfortunately, we failed to find any use for such marking in our further analysis, but if one wants to use our code on Github, we think s/he should be aware of such feature.*

        2.3.3. If they both form a clade, or they both do not form a clade, the hole remains unclassified and is not analyzed. It is, however, recorded as a “strange example” in the script log.

2.4. The target hole is removed from the holes list, and a new iteration through paragraphs 2.1-2.4 is initiated.

3. After all the insertions and deletions are classified, species undergo filtering. Species with genes (1) with nucleotide number not divisible by 3 or (2) without a start and/or an end or (3) having an inner stop are removed. *The reason this step is not performed at the start is that species with such unreliable and possibly erroneously sequenced genes could still help us to determine insertions and deletions in species with trustworthy genes.*

4. For every species in the alignment that is left (hereafter, target species), paragraphs 4.1-4.6 are executed (insertions and deletions of that species are or are not classified as pCFM).

4.1. A list of insertions and deletions (indels) this species carries is formed. Indels of length divisible by 3 are dropped from this list (i.e. further steps assume they never existed). *This step is performed because such indels are not of any interest in the search for pCFMs, and the number of pCFMs candidates will matter in paragraph 4.3.*

4.2. Indels, which are long (>20 nucleotides) and common for target species and all the descendants of this species’s parent node are also dropped from the list. *This is a mechanism for not considering long unaligned regions in basal species to be indicators of insertions in other species (suppose, for example, a hole of length 100 in a couple of basal species, which is most probably a defect of exon alignment. The algorithm will define sequences in all other species as insertions, which would not be appropriate. This procedure is taken in order to get rid of such abnormal “insertions”). This step is not crucial for general understanding of an algorithm, but if one wants to use our code on GitHub, we think s/he should be aware of that feature as well.*

4.3. If more than 2 indels are left in the list, the species is dropped (i.e. paragraphs 4.4-4.5 are not executed). *The reasoning behind this step is that such species are likely to be somewhat ill-sequenced or ill-aligned. However, handling species with multiple pCFMs was implemented, but wasn’t used (the parameter* brutal_conditions *in the script results in bypassing this step).*

4.4. The indels are classified as pCFM if one of two conditions are met: (1) indels have the same name (insertion-insertion or deletion-deletion) and sum of their lengths is divisible by 3 or (2) indels have different names (insertion-deletion or deletion-insertion) and the difference of their lengths is divisible by 3.

4.5. If pCFMs are detected, previously dropped species with the same indels (paragraph 3) are added into consideration. *The reasoning here is that if they carry the same indels as trustworthy species, they are probably also trustworthy. This step is performed to gain more support in our analysis: we consider the cases where multiple species carry the pCFM to be more reliable.*

4.6. If pCFMs are detected, it is checked, if both frameshfting indels from the pair happened simultaneously or not. For that two last common ancestors are compared: the last common ancestor of the species carrying the first frameshifting mutation of the pair and the last common ancestor of the species carrying the second. If these common ancestors are the same, frameshifting mutations happened simultaneously, else they did not.

5. After the iteration through all the species, for each pCFM the following information is added to the output : species in which pCFM was found, the names of mutations it is comprised of (insertion/deletion), lengths, positions, and simultaneity of their happening. An output is given in a form of two tsv tables: one for the simultaneous mutations, another for non-simultaneous.

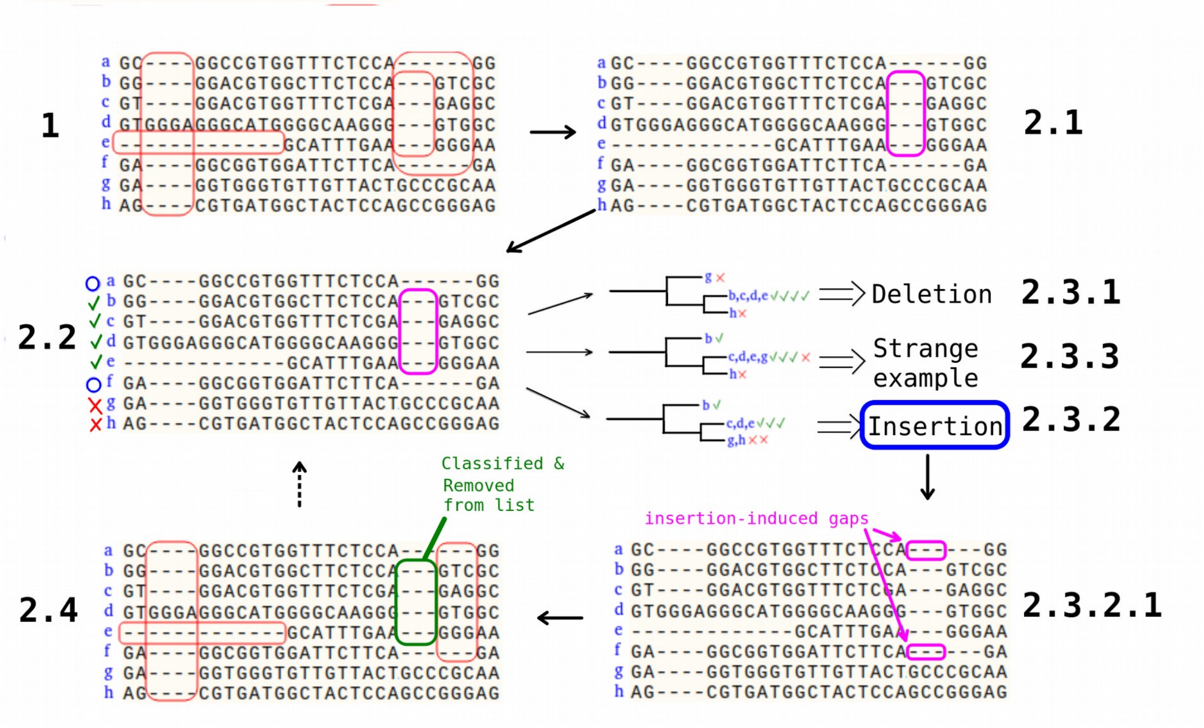


Supplementary Figure 1. A schematic representation of the first part of the algorithm of search for indels in the alignment: classification of indels. A fragment of an alignment is shown, with letters a-h denoting different species. Red frames correspond to distinct holes. Green check marks flag species sharing the considered hole, and red crosses, those not sharing it. Blue circles flag the rest of the species (for which the presence of the hole is not unequivocal). The numbers correspond to the paragraphs in the algorithm description.

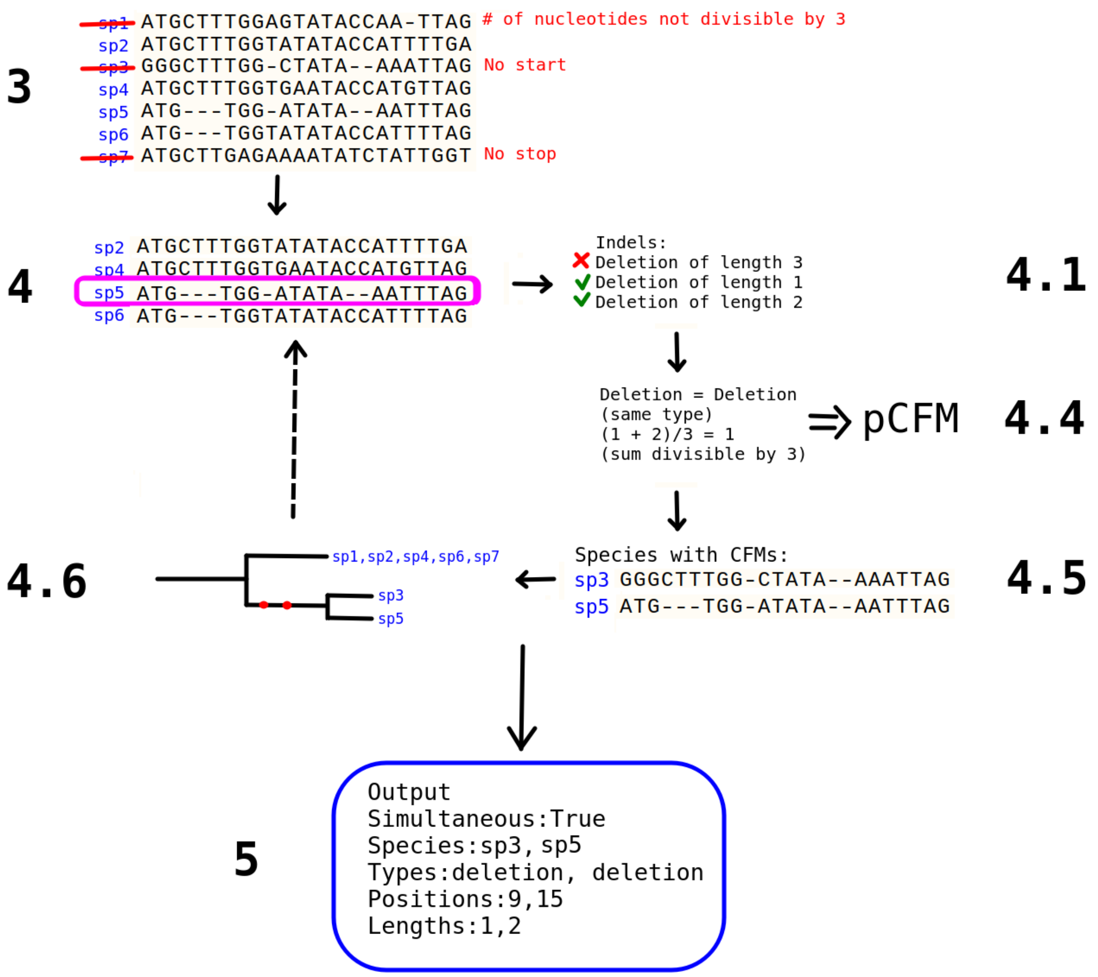


Supplementary Figure 2. A schematic representation of the second part of the algorithm of search for indels in the alignment: filtering and identification of pCFM. Species are denoted as sp1-sp7. The numbers correspond to the paragraphs in the algorithm description.

### Supplementary Table 1

Supplementary Table 1. The list of species in which the pCFMs were detected for each of the 11 high-confidence pCFM-carrying genes.

| Gene | Species carrying the pCFM |
|---|---|
| *RAB36* | calJac3, papHam1, panTro4, chlSab1, saiBol1, rheMac3, gorGor3, ponAbe2, hg19, nomLeu3, macFas5 |
| *ARHGAP6* | panTro4, macFas5, papHam1, ponAbe2, nomLeu3, hg19, rheMac3, chlSab1, gorGor3 |
| *INSL6* | vicPac2, bosTau7, susScr3, turTru2, panHod1, oviAri3, camFer1, orcOrc1, capHir1 |
| *NCR3LG1* | ponAbe2, panTro4, hg19, gorGor3 |
| *SPATA24* | cavPor3 + *Cavia aperea* (ENSEMBL) |
| *RASSF4* | myoDav1, eptFus1, myoLuc2 |
| *Cyp6a23* | Bactrocera_tryoni, Bactrocera_oleae, Bactrocera_latifrons, Bactrocera_dorsalis |
| *Diap1* | droMir2, D_athabasca, droPse3, droPer1 |
| *Osi10* | droMoj3, D_navojoa, D_hydei, D_arizonae |
| *wds* | D_serrata, droKik2 |
| *CG31530-RA* | droTak2, droEug2, dm6, droSec1, droEle2, droKik2, droYak3, D_serrata, droSim2, droBia2, droSuz1 |

#### Supplementary Table 2

Supplementary Table 2. Support from NCBI and Uniprot databases obtained for each of the 11 pCFM-carrying genes. The columns "evidence" and "source" indicate the type of evidence (protein or mRNA or predicted gene) and the database this evidence was obtained from. In the last column, the ID for the corresponding database (RefSeq ID or UniProt ID) is presented. For each gene and for each of the variants (with and without the pCFM), the evidence of type "mRNA" or "protein" is listed for all species for which such evidence is available. The evidence of the type "predicted gene" is listed only if no "mRNA" or "protein"-level evidence is available, and just for one of the species.

| Gene | Species | Evidence | Source | ID |
|---|---|---|---|---|
| *RAB36* | **With pCFM: hg19** | mRNA | NCBI | NM_004914.4 |
| | Without pCFM: mm10 | mRNA | NCBI | NM_001359270.1 |
| *ARHGAP6* | **With pCFM: hg19** | mRNA | NCBI | NM_013427.2 |
| | | protein | UniProt | O43182 |
| | **With pCFM: panTro4** | mRNA | UniProt | H2QYA3 |
| | Without pCFM: mm10 | mRNA | NCBI | NM_009707.4 |
| *INSL6* | **With pCFM: bosTau7** | mRNA | NCBI | NM_001077521.2 |
| | Without pCFM: hg19 | mRNA | NCBI | NM_007179.3 |
| | Without pCFM: mm10 | mRNA | NCBI | NM_013754.1 |
| | Without pCFM: rn5 | mRNA | NCBI | NM_022583.1 |
| *NCR3LG1* | **With pCFM:** | mRNA | NCBI | NM_001202439.2 |

| | | | | |
|---|---|---|---|---|
| | **hg19** | protein | UniProt | Q68D85 |
| | Without pCFM: nomLeu3 | predicted gene | NCBI | XM_003254281.4 |
| | | mRNA | UniProt | A0A2I3FX64 |
| *SPATA24* | **With pCFM: cavPor3** | predicted gene | NCBI | XP_003477496.1 |
| | Without pCFM: mm10 | protein | UniProt | Q6P926 |
| *RASSF4* | **With pCFM: myoDav1** | predicted gene | NCBI | XM_006761419.2 |
| | Without pCFM: eriEur2 | predicted gene | NCBI | XM_007516816.2 |
| *Cyp6a23* | **With pCFM: Bactrocera_latifrons** | predicted gene | NCBI | XM_018936230.1 |
| | Without pCFM: dm6 | mRNA | UniProt | Q9V771 |
| *Diap1* | **With pCFM: droPer1** | predicted gene | NCBI | XM_026989947.1 |
| | Without pCFM: dm6 | protein | UniProt | Q2430 |
| *Osi10* | **With pCFM: droMoj3** | predicted gene | NCBI | XM_001999204.2 |
| | Without pCFM: DroVir3 | predicted gene | NCBI | XM_002058517.2 |
| *wds* | **With pCFM: Dserrata** | predicted gene | NCBI | XM_020947031.1 |

| | | | | |
|---|---|---|---|---|
| | Without pCFM: droAna3 | predicted gene | NCBI | XM_001966487.3 |
| *CG31530-RA* | **With pCFM: droKik2** | predicted gene | NCBI | XM_017175617.1 |
| | Without pCFM: dm6 | mRNA | UniProt | D2NUL5 |

### Supplementary Figure 3

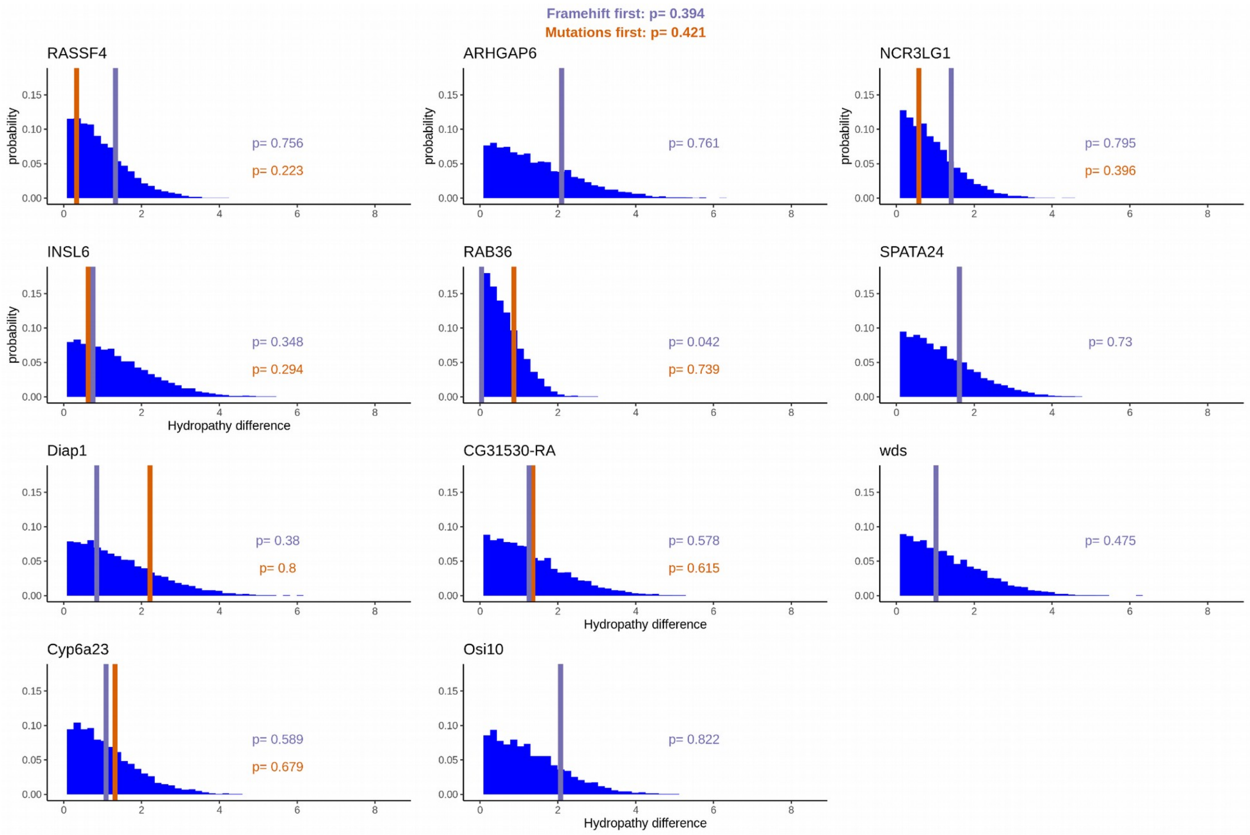


Supplementary Figure 3. The effect of pCFM on hydropathy of the encoded amino acid sequence. The notation is the same as in Figure 5.

### Supplementary Figure 4

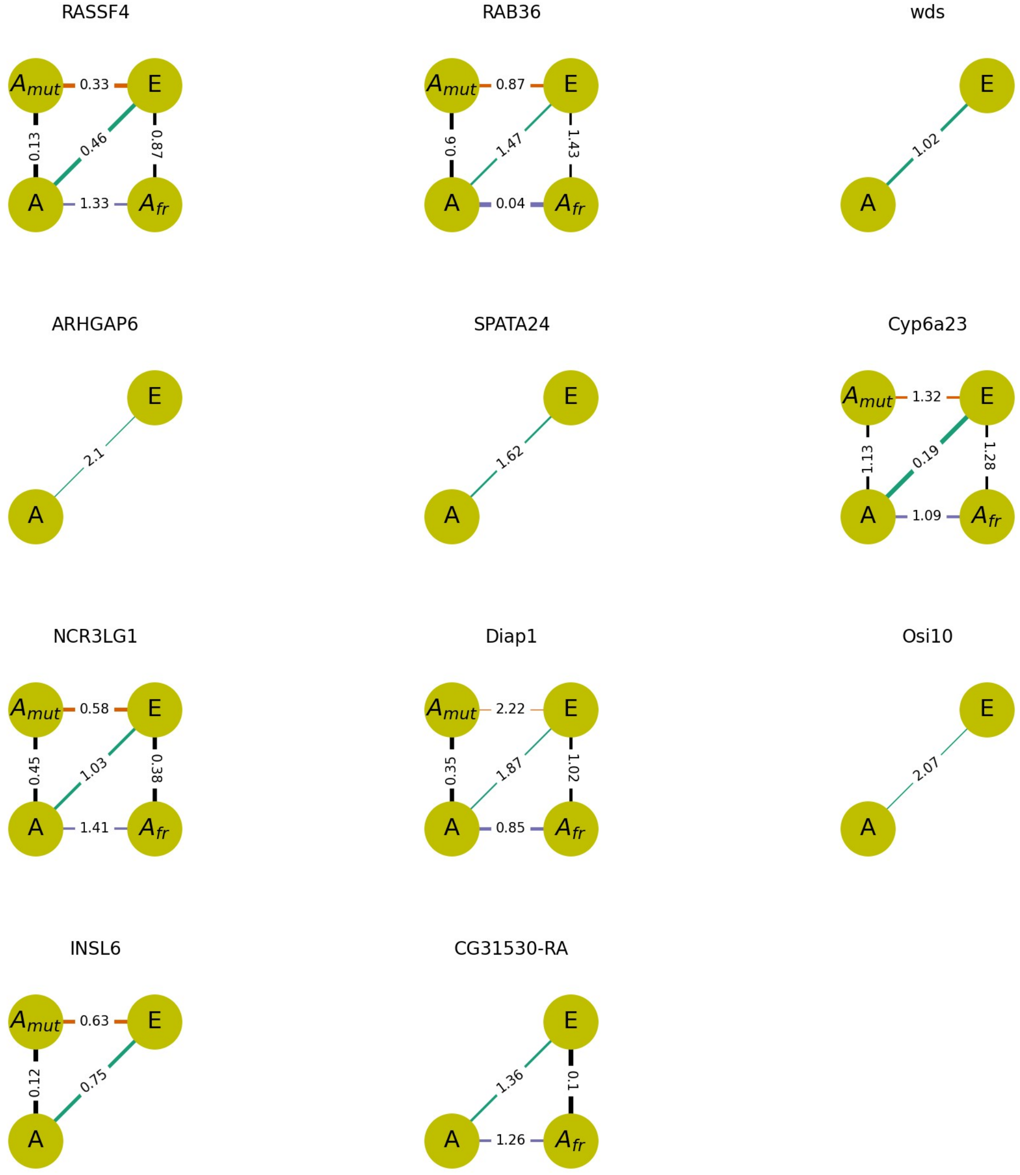


Supplementary Figure 4. Effects of pCFM and amino acid substitutions occurring on the same phylogenetic branch as pCFM on the physico-chemical properties of the encoded protein, according to the difference in hydropathy of the encoded amino acid sequence. The notation is the same as in Figure 6.

### Supplementary Figure 5

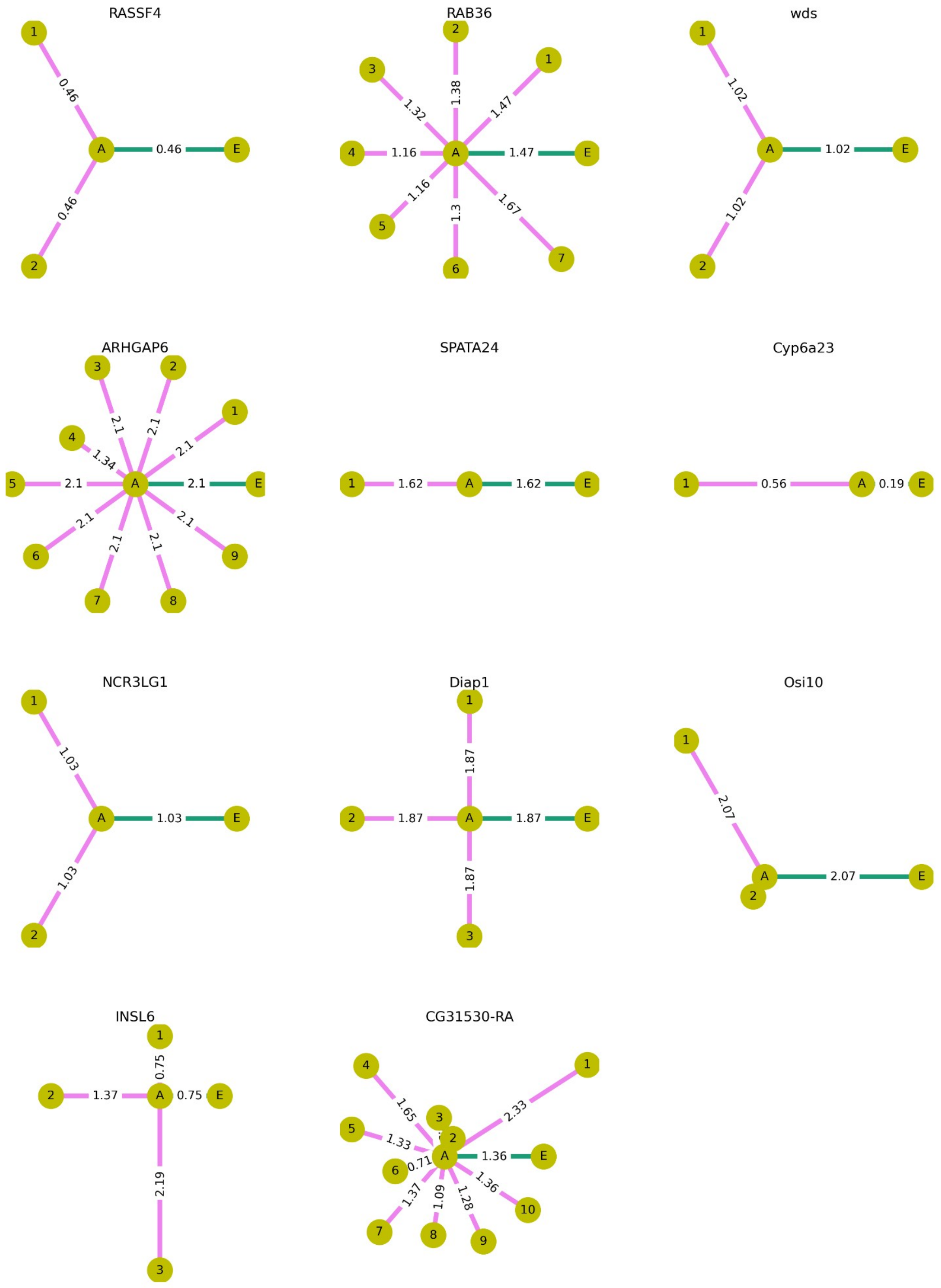


Supplementary Figure 5. Effects of amino acid substitutions occurring on the phylogenetic branches descendant to pCFM-branch on the physico-chemical properties of the encoded protein, according to the difference in hydropathy of the encoded amino acid sequence.The notation is as in Figure 7.
